# Similarity identification in gene expression patterns as a new approach in phenotype classification

**DOI:** 10.1101/110130

**Authors:** Seyed Ali Madani Tonekaboni, Venkata Satya Kumar Manem, Nehme El-Hachem, Benjamin Haibe-Kains

## Abstract

Stratifying healthy and malignant phenotypes and identifying their biological states using high-throughput molecular data has been the focus of many computational approaches during the last decade. Using multivariate changes in expression of genes within biological pathways, as fingerprints of complex phenotypes, we developed a new methodology for Similarity Identification in Gene expressioN (SIGN). In this approach, we use centroid classifier to identify phenotype of each biological sample. To obtain similarity of a given biological sample with classes of phenotypes, we defined a new distance measure, transcriptional similarity coefficient (TSC) which captures similarity of gene expression patterns between a biological pathway in two samples or populations. We showed that TSC, as an interpretable and stable distance measure in SIGN, captures all oncogenic hallmarks for breast cancer even with low sample size, by comparing healthy and patient tumor samples in the largest breast cancer dataset. In this study, we demonstrate that SIGN is a flexible, yet robust approach for classification based on transcriptomics data. Comparing early and late relapses within each molecular subtypes of breast cancer, our method enabled subtype-specific stratification of breast cancer patients into groups with significantly different survival. Moreover, we used SIGN to classify with more than 99% specificity the site of extraction of healthy and tumor samples from the Genotype-Tissue Expression (GTEx) and The Cancer Genome Atlas (TCGA) datasets. We showed that SIGN also enables robust identification of hematopoietic stem cell and progenitors within the hematopoietic hierarchy. We further explored chemical perturbation data in the Connectivity Map (CMAP) database and showed that SIGN was able to classify seven classes of drugs based on their mechanism of action. In conclusion, we showed that SIGN can be used to achieve interpretable and robust transcriptomic-based classification of healthy and malignant samples, as well as drugs based on their known mechanism of action, supporting the generalizability and relevance of the method for the analysis of gene expression profiles.

## INTRODUCTION

Messenger RNA expression is an important feature representative of the biological state of a cellular population. Activity of tissue-specific genes, master regulatory transcription factors, onco- and tumor suppressor genes all can play important roles in variety of healthy and disease phenotypes [1–3]. For example, cancer is a genetic disease and gene expression is a reflection of various carcinogenic features in a tumor. External stress such as drug treatment or hypoxia as well as other microenvironmental conditions of a tissue could potentially affect the mRNA transcription [4,5]. Furthermore, the spatial and temporal dynamics of these conditions also dictate the functional behaviour of the phenotype in a system [6].For instance, genes expressed in tumor epithelium and stromal compartments, have been shown to have different transcriptomic profiles, which are significantly associated with clinical outcome of breast cancer patients [7].

There have been tremendous efforts during the last fifteen years in developing computational approaches to compare expression of genes in different biological conditions and subsequently assign their contribution into known biological pathways [8]. The three main categories of the methodologies, including singular enrichment analysis (SEA), gene set enrichment analysis (GSEA), and modular enrichment analysis (MEA) [8], rely on summarizing the data as a list of genes or a vector of fold change in expression of genes between two populations (Fig. 1). Hence, the methods will loose the biological information by dismissing covariance of gene expression in the samples within each population. Hence, these methods disregard the dependency of changes in expression of genes in biological pathways.

**Figure 1.**
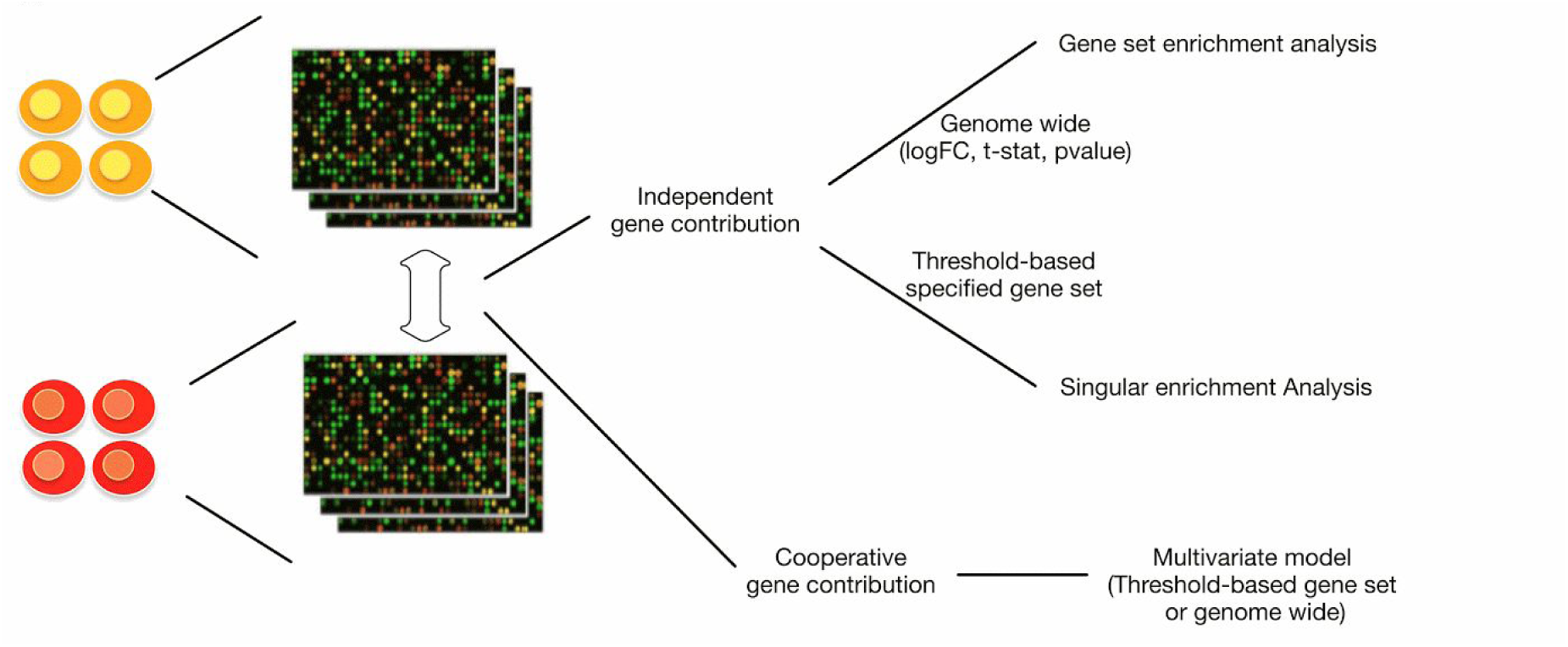
Categories of methods developed to identify genes or pathways with different expression value or pattern between two populations (red and orange cells in this figure are considered as schematic representation of two populations).

Alternatively there have been multivariate models developed to capture variations between two populations or obtain gene expression based biomarkers of a phenotype [9]. For example, a great deal of research effort has been directed towards building gene expression based biomarkers, called as gene signatures to identify highly proliferative tumors, metastatic tumors, prognosis, responders and nonresponder using this approach [10–13]. However, multivariate models are susceptible to other issues such as risk of overfitting as there is huge difference in accuracy of these models in training dataset for learning and validation dataset [14].In addition, considering feature selection as an NP-hard problem, there is no unique and globally optimal solution while the available methods and studies report a unique signature [15,16] in spite of existence of multiple signatures with equivalent discriminative power. Moreover, it is difficult to interpret the signatures difficult to interpret because known biological pathways have not been considered during the discovery/fitting process.

To circumvent the above limitations associated with enrichment-based and multivariate-based methodologies, we sought to develop a new statistic to consider cooperative contribution of genes within biological pathways as an interpretable approach which captures variability of expression of genes within biological samples. To compare the list of pathway representative of each population, we defined transcriptional similarity coefficient (TSC). Using TSC, we captured all oncogenic hallmarks with TSC for different subtypes of breast cancer without being limited to choice of tumor samples, imbalance in normal and tumor sample populations, and number of samples in subtype-specific tumor populations.

Upon validating performance and stability of TSC, we used it as a similarity measure in a nearest centroid classification framework called Similarity Identification in Gene expressioN (SIGN) to identify phenotypes under normal and stress conditions. We primarily test the method for identification of normal and cancerous samples and further explored its performance in stratifying breast cancer patients based on their survival. We proved its significant performance in breast cancer patient stratification. To demonstrate the versatility of SIGN in identifying phenotypes in different biological states, we implemented it to classify healthy and malignant patient and patient-derived xenograft samples. Moreover, we illustrated ability of SIGN in detecting type of chemotherapy that cancer cells have been exposed, and caused changes in gene expression patterns in their biological pathways, using chemical perturbation data in connectivity mapping database [17].

## METHODS

We hypothesized that gene expression patterns within biological pathways are specific to each phenotype. To investigate validity of this hypothesis, we defined transcriptional similarity coefficient (TSC) which captures relative changes of gene expression within biological pathways between two biological samples. In this regard, we primarily considered each phenotype as its biological pathways including biological processes (BP), molecular functions (MF), and cellular components (CC) as defined in the Gene Ontology [18](Fig. 2A).

**Figure 2.**
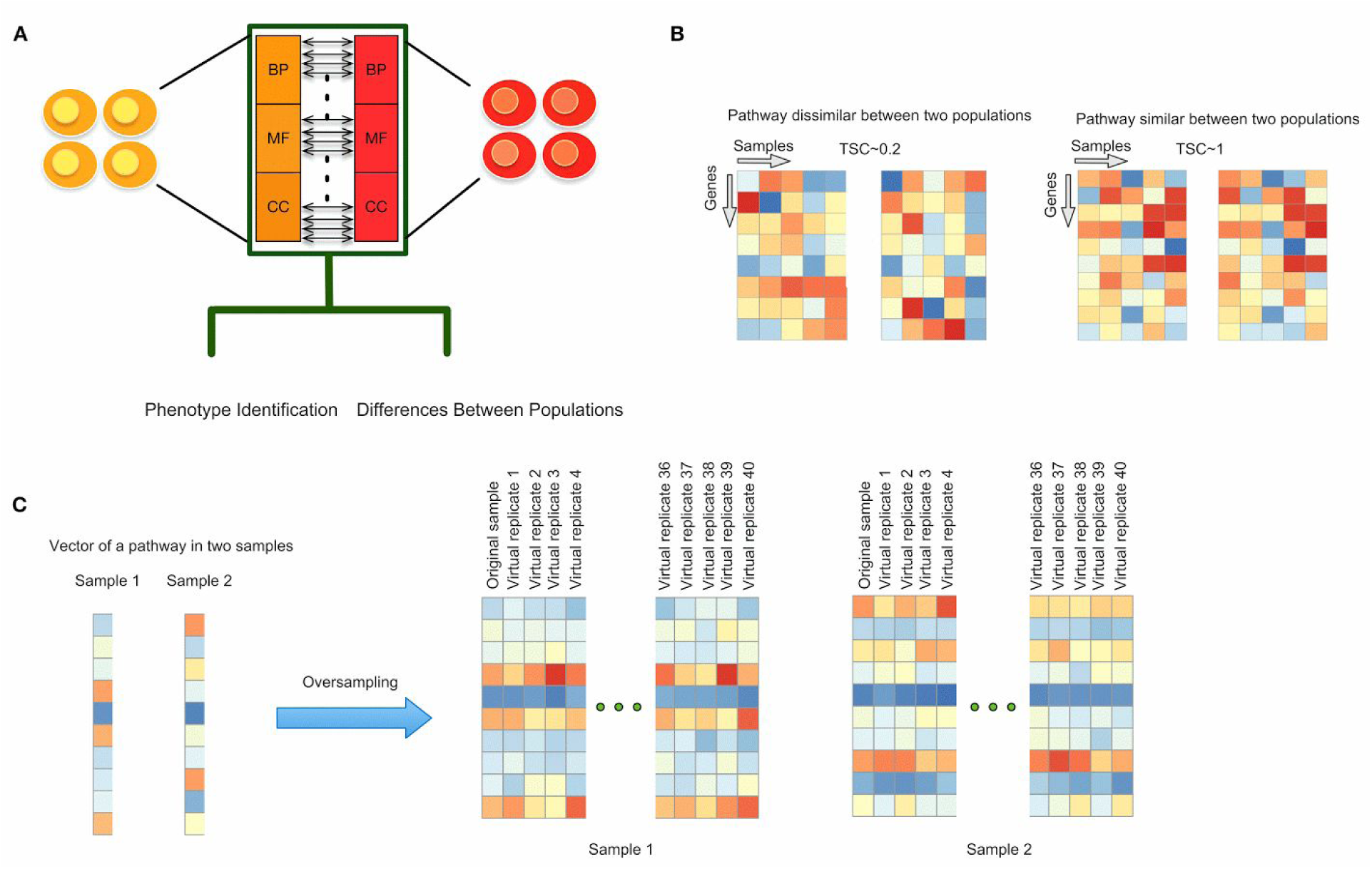
Transcriptional similarity coefficient (TSC) as a new statistic to identify genes or pathways with different expression value or pattern between two populations (red and orange cells in this figure are considered as schematic representation of two populations). **A**) Illustrating each biological sample as its biological processes (BP), molecular functions (MF), and cellular components (CC) as have been considered in gene ontology (GO) terms; **B**) Transcriptional similarity coefficient (TSC) values and for high and low similarity of transcriptional patterns; **C**).

### Transcriptional similarity coefficient

Let *P* be the matrix of expression of genes within a pathway for one set of biological samples where rows are genes and columns are samples. Afterward, we obtain transcriptional similarity coefficient (*TSC*) between the two matrices using modified RV-coefficient [19] as follows

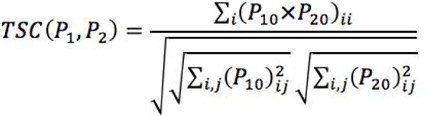

where *P*_*1*_ and *P*_*2*_ represent the matrix of gene expressions of a given pathway in two set of samples (population 1 and 2), *i* represents one row (i.e., one gene) within each matrix, *j* represents one column (i.e., one sample) within each matrix, and P_m0_

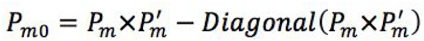

where the *Diagonal* function sets to zero the elements of the matrix that are not in the diagonal. The range of the TSC score lies in [−1,1]. Higher scores will be equivalent to higher similarity of gene expression pattern of a given pathway between two populations (Fig. 2B). Therefore, *TSC* represents the similarity of a given pathway between two samples and/or populations. We identify dissimilar pathways between two populations relying on distribution of TSCs between the populations. We consider *i*th pathway to be dissimilar if *T SC*_*i*_ − *median*(*Q*(*T SC*)))/*mad*(*Q*(*T SC*)) is less than −1.69.

To capture gene expression patterns for a single sample, additional virtual replicates are built by adding uniform noise to expression of each gene (Fig. 2C), similar to the concept of oversampling [20].Hence, instead of a single sample, we will have one original gene expression profile and a series of virtual samples which helps us to build a matrix of gene expression pattern for the single sample.

### Phenotype identification

Upon validating TSC as a coefficient capturing biological difference between phenotypes, we developed similarity identification for gene expression (SIGN) as an approach to classify phenotypes based on their transcriptional pattern similarity. In this method, expression pattern in GO terms is used as fingerprint of a given phenotype.

To classify a single sample, we use centroid classifier to obtain the most similar class of phenotype to a given biological sample (Fig. 3). We used TSC as a measure of similarity of biological pathways between the given sample and the classes. Defining *Q*(*T SC*) as distribution of *TSC* between all pathways in the given sample and each phenotypic class, we defined

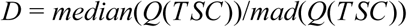

**Figure 3.**
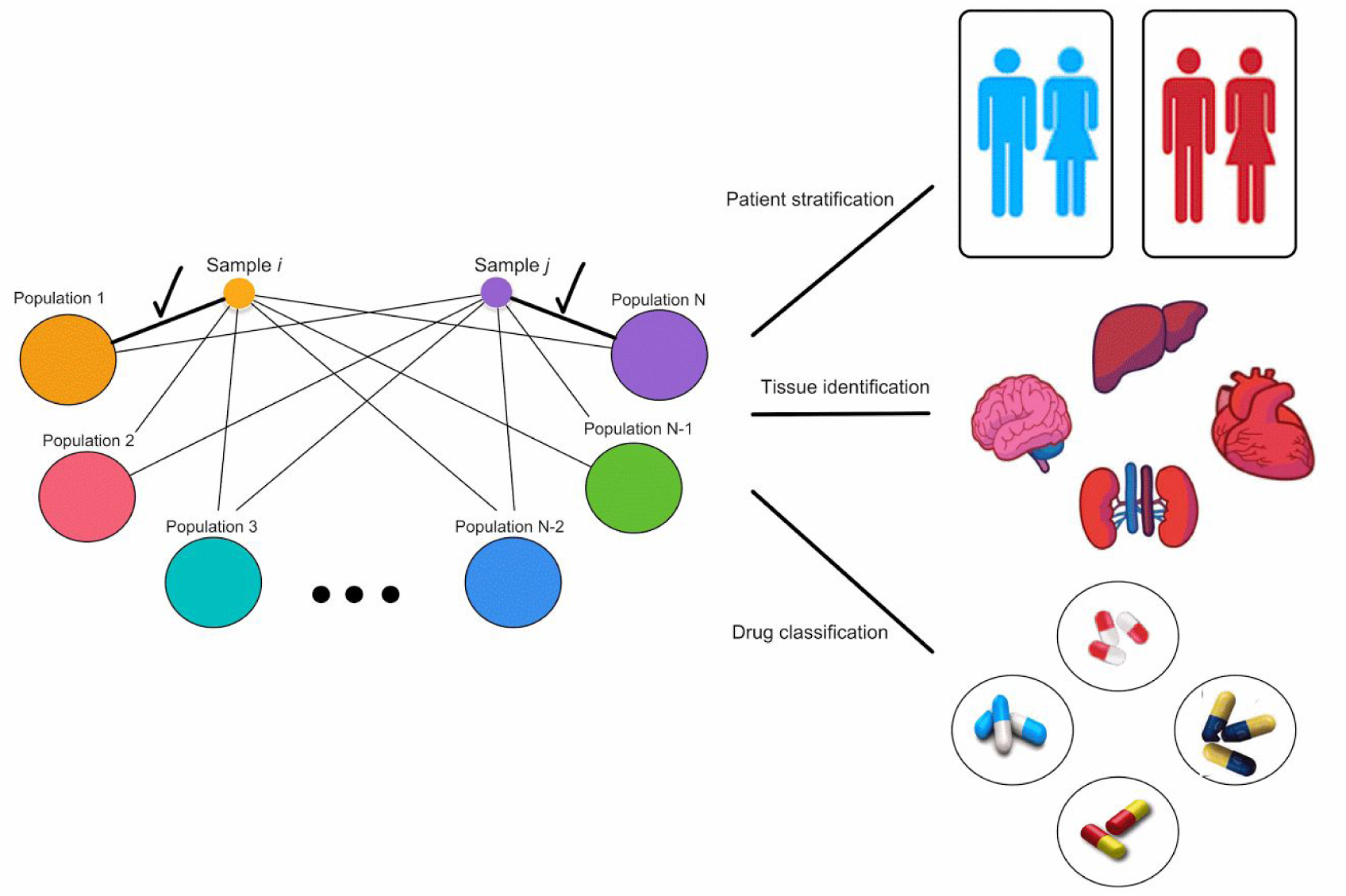
SIGN classification method and its application. Using similarities between samples and reference population to identify the population with the nearest centroid as the main part of SIGN method; and its application in stratifying patients based on their survival rate, identifying site of extraction of tissue samples, and classifying drugs based on their mechanism of action.

Where *D* is the distance measure between the given sample and each class. We then find the population with the nearest centroid to the given sample and consider the phenotype of that population as the phenotype of the examined sample. Moreover, we defined Delta as the difference between similarity, defined as *median*(*D*(*T SC*))/*mad*(*D*(*T SC*)), of a biological sample between two phenotypic classes.

### Mapping pathways to hallmark gene sets

To have a summary dissimilar pathways between healthy and malignant population of samples, we map list of genes within dissimilar pathways between the populations and map them on the hallmark gene sets provided in MSIGDB [21].Implementing 1,000 permutation test, by randomly selecting same number of genes from all the genes provided in molecular profiles of the samples, we identify significantly enriched hallmarks being dissimilar between the populations.

### Breast cancer molecular subtyping

We used the SCMOD2 model [22] implemented in the *genefu* package [23] to assign each tumor samples in Molecular Taxonomy of Breast Cancer International Consortium (METABRIC) database of breast tumors into the four established molecular subtypes of breast cancer: ER-/HER2-, HER2+, ER+/HER2- low and high proliferation.

### Survival Analysis

We used the METABRIC dataset (Supplementary Table 1) [24], including 1992 breast tumor samples, and overall survival as an endpoint. We defined and used it to *Delta* = (*similarity to good survival cohort*) − (*similarity to poor survival cohort*) and used it to identify difference of similarity of each patient tumor sample to good and poor survival populations. We assessed the prognostic value of *Delta* as a continuous value using the D index and as a discrete value using the logrank test for Kaplan-Meier survival curves as implemented in the *survcomp* R package [25]. We split the population cohort patients, for each subtype, into two groups of low, intermediate, and high survival to include ~10% of the patients, in each subtype cohort, in low and high survival categories and 80% in the intermediate category.

### Similarity of pathways based on genomic perturbation data

We used the *PharmacoGx* package to extract drug perturbation signatures in connectivity mapping (CMap) database [17]. Briefly, given a PharmacoSet of the perturbation experiment type, and a list of drugs, a drug-specific signature for the effect of chemical concentration on the molecular profile of a cell will be computed using a regression model adjusted for experimental batch effects, cell specific differences, and duration of experiment. For each gene, we used the t-statistic of the drug concentration coefficient to characterize the drug effect. We subsequently build matrices of t-scores of genes within every biological pathways for one set of chemical perturbations. We used TSC to obtain similarity of these matrices and assessed similarity of the effect of two drugs on biological pathways considering distribution of TSCs for all pathways.

## RESULTS

The available statistics in literature for comparison of pathways, based on expression of their included genes, between two populations rely on enrichment of their genes as being up- or down-regulated in one population versus another. Hence, we need a new statistic to investigate our hypothesis in which relative changes in expression of genes within biological pathways (i.e. gene expression patterns; Fig. 2B) is fingerprint of a phenotype. In this regard, we defined a new coefficient called transcriptional similarity coefficient (TSC). We consider each pathway as a matrix in which the rows represents the genes within the pathways and columns are the samples in a given population (Fig. 2B). Hence, TSC estimates the similarity of gene expression patterns for the same pathway in two populations. In this study we leveraged TSC to compare healthy and tumor samples at the pathway level, predict patient survival, classify samples based on their tissue of extraction, identify subpopulations within the hematopoietic hierarchy and classify drugs based on their transcriptomic effects on cancer cell lines.

### Hallmarks of cancer

To identify oncogenic pathways, we used TSC to compare the gene expression patterns of pathways across a large cohort of breast tumor and healthy samples in METABRIC dataset [24]. We compared healthy samples with all tumor samples as well as subtype-specific patient cohorts to capture subtype-specific oncogenic pathways. Using TSC, we identified all hallmarks of cancer, reported as hallmark gene set in MSigDB [21], to be oncogenic in at least one subtype of breast cancer (Supplementary File 1). There were four hallmarks which were oncogenic in subtype-specific manner, and not significant for all the tumor samples together, including DNA Repair, Unfolded Protein Response, MYC Targets, and Oxidative Phosphorylation.

We identified DNA Repair as hallmark of TNBC subtype of breast cancer. It could be due to deficiency of this subtype of breast cancer, with respect to other subtypes, in homology-dependent repair for the repair of double strand breaks [26,27]. Oxidative phosphorylation which was another subtype-specific identified hallmark contribute to the survival and growth of the cancer cells [28]. It is well known that tumor epithelial cells induce glycolysis in fibroblasts, which in turn secrete lactate and pyruvate metabolites along with other growth factors. As TNBC and HER2 are representative of highly aggressive tumors with proliferative activity, oxidative phosphorylation of cancer cell metabolism could be expected [28].Moreover, Unfolded Protein Response is identified by TSC as being specific oncogenic hallmark for ER-positive and triple negative breast cancer subtypes as being reported as causes of resistance to antiestrogens and chemotherapeutics in these two subtypes [29]. Although MYC hallmark gene set is identified as ER-positive specific oncogenic hallmark, we identified targets of MYC, as a key oncogene [30], such as Epithelial to Mesenchymal Transition, Angiogenesis, and Apoptosis being oncogenic for all subtypes of breast cancer.

The oncogenic pathways identified for breast cancer subtypes by comparing healthy and tumor samples in METABRIC by TSC showed us the validity of this statistic as a phenotype specific statistic of pathway activity. We sought stability of the results by bootstrapping tumor samples in each subtype-specific patient cohorts. We randomly chose 30% of the samples 100 times and compared the gene expression patterns in biological pathways using TSC between the obtained population and healthy breast samples in METABRIC. Out of 50 hallmarks, only 3 of the pathways showed instability which had different significance in more than 5% of the sampling repetitions for TNBC subtype (Supplementary File 2). The instability in other subtypes was lower as 2 pathways were unstable for HER2 and ER+/HER2- high proliferation subtypes and 1 for ER+/HER2- low proliferation subtype (Supplementary File 2). Hence, TSC not only capture the pathways phenotype specific pathways but also it does not depend on the samples one may use to investigate specificity of gene expression pattern in a phenotype of interest.

Moreover, we explored the effect of imbalance sample number and low sample number by repeating the bootstrapping with low tumor sample number. In this case, we randomly chose 10 samples 100 times and compared the gene expression patterns in biological pathways using TSC between the obtained population and 144 healthy breast samples in METABRIC. In spite of high level of imbalance (10 versus 144) and low tumor sample number, there were at most 6 hallmarks which had instability in significance in more than 5% of the sampling repetitions which was for TNBC subtypes. There were 3, 4, and 5 unstable hallmarks for HER2, ER+/HER2- high proliferation, and ER+/HER2- low proliferation subtypes, respectively.

Upon validation of performance of TSC in identifying hallmarks of breast cancer subtypes and its stability with respect to variation in patient samples and using low sample number and being robust using imbalanced populations, we hypothesized that TSC is a robust measure of gene expression pattern similarity between phenotypes. Hence, we sought to use this measure to classify phenotypes. We developed a classification approach, similarity identification in gene expression (SIGN) using TSC as a measure of similarity of biological pathways. In this approach, we consider each sample as a node in a network and find a population which has the nearest centroid to a given sample and consider it as its phenotype. In this regard, we obtain TSC for transcriptional similarity of biological pathway between the sample and each population and use median over mad (absolute deviation from data median) of TSCs as the distance measure between the given sample and each population (Fig. 3).

We used SIGN to identify healthy and tumor samples in METABRIC by investigating if healthy or tumor population has the most similarity to a given sample based on transcriptional patterns of biological pathways. We identified 1933 out of 1992 tumor samples and 142 out of 144 healthy samples in METABRIC (Matthew Correlation Coefficient [MCC]=0.82, p<0.001). Hence, we hypothesized that gene expression patterns within biological pathways can be fingerprints of phenotypes and representative of their biological state. Therefore, we further explored the ability of SIGN in stratifying patient tumors based on their survival rate as a more challenging question.

### Patient stratification based on their survival

We used the METABRIC dataset and overall survival as an endpoint and examined ability of SIGN in stratifying patients based on their survival. We defined poor and good survival groups as the patients who die the earliest after diagnosis and the patients who survived the longest (10% of the cohort in each category). We used poor and good survival patient cohorts, each including ~10% of the population, to train the model and check validity of Delta in stratifying patients. Hence, we considered the rest of the population, patients with intermediate survival, as test set to assess performance of SIGN in patient stratification. For each patient, the difference (referred to as *Delta*) between the TSC score of the poor and good groups is computed so that patients with positive Delta should have higher survival than the patients with negative *Delta*. We assessed the prognostic value of Delta using the concordance index for each molecular subtype separately (Fig. 4). SIGN yielded significant prognostic value for all the subtypes (concordance indices > 0.7, p<0.002; Fig. 4A), with estrogen receptor (ER)-negative/human epidermal growth factor 2 (HER2)-negative and the HER2+ breast cancer being the most challenging to classify (concordance index=0.72 and 0.73, for ER-/HER2- and HER2+, respectively) while ER+/HER2- low proliferation have the highest concordance index (0.82). To further visualize the prognostic value of SIGN, we computed the Kaplan-Meier survival curves based on the sign of Delta predictions for the patients with intermediate survival (logrank p<0.05; Fig. 4B).

**Figure 4.**
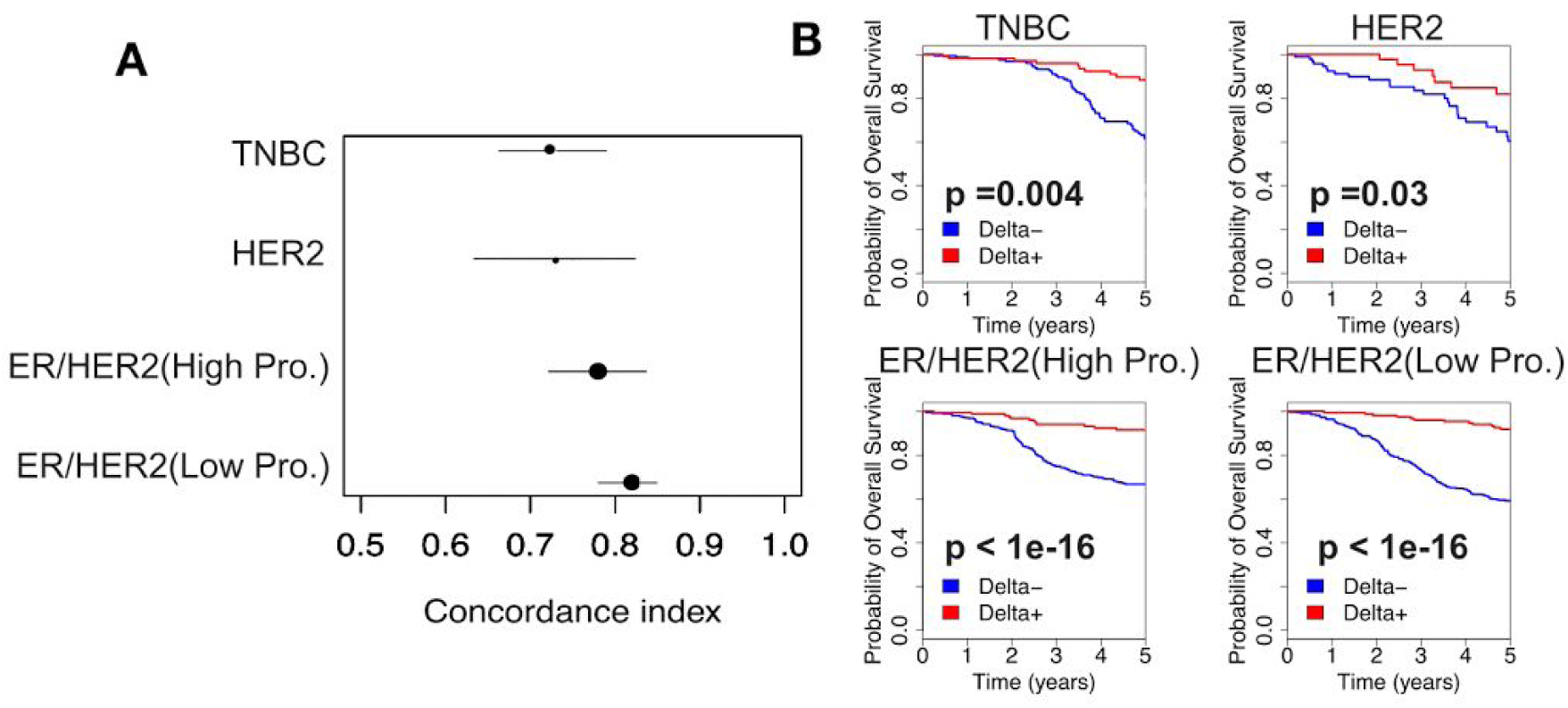
Subtype-specific survival analysis using SIGN. **(A)** Forest plot reporting the concordance indices (plain circles) along with the 95% confidence interval (horizontal bars) for stratification of patients with intermediate survival in the METABRIC dataset. (**B**) Kaplan-Meier 5-year survival plot along with the logrank test of intermediate survival patient stratification using SIGN. Intermediate survival cohorts for ER-/HER2-, HER2+, ER+/HER2- are those patients who survived in the range 1.5-12, 1.5-14, and 3-15 years, respectively.

### Tissue type identification

Being successful in stratifying breast cancer patients as different phenotypes from same tissue, we sought to identify gene expression pattern similarities between different healthy and malignant tissue types. We implemented SIGN to identify the site of extraction of each tissue sample in the Genotype-Tissue Expression (GTEx) dataset based on their gene expression profiles [31] (Fig. 5A). Using leave-one-out cross validation strategy, average balanced accuracy (½*(TP/P+TN/N)) of the predictions were 99.4% for the 13 tissues with more than 70 samples (Supplementary Table 2). SIGN could identify samples of blood, lung, and muscle tissues with 100% sensitivity (Fig. 3A).

**Figure 5.**
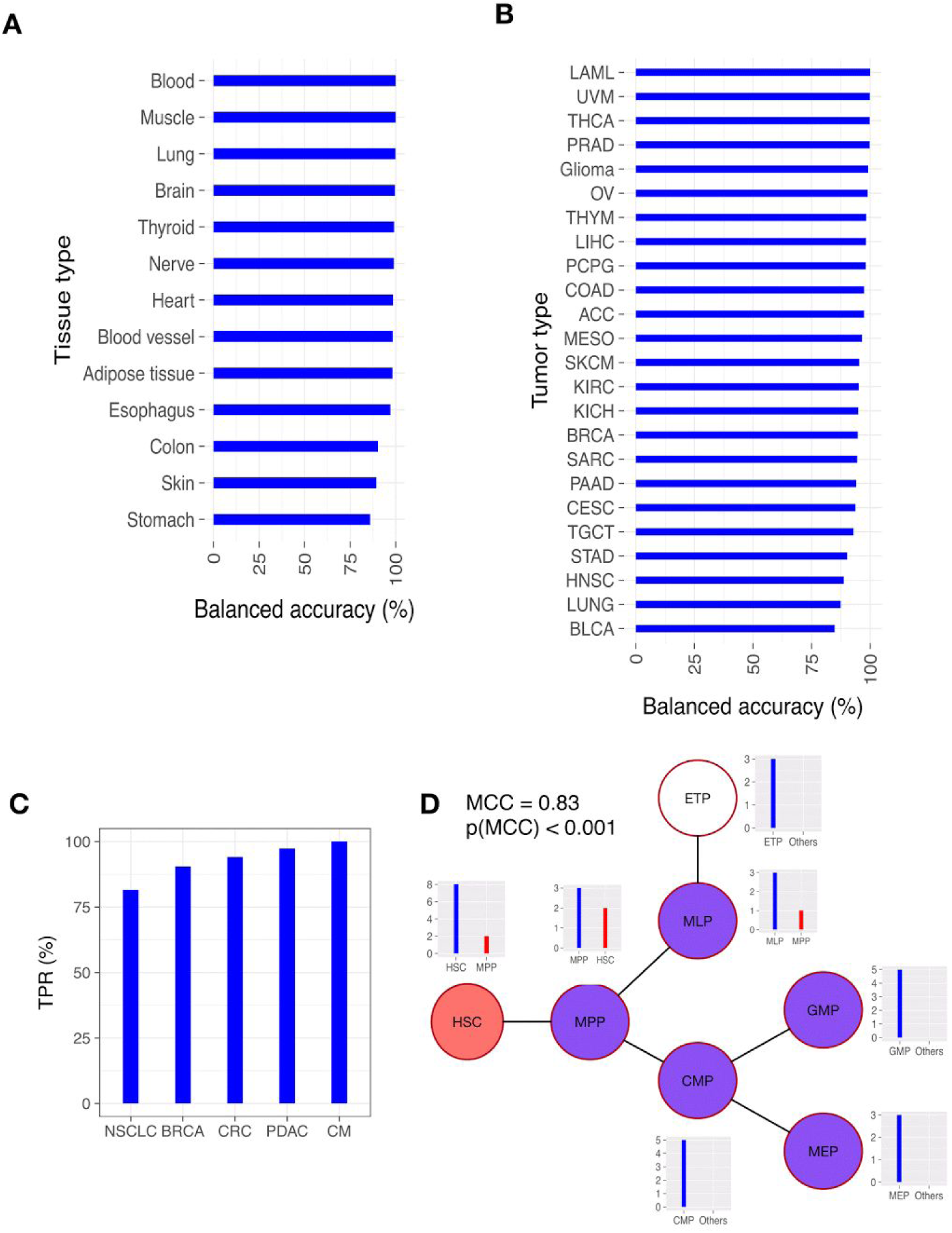
Identification of site of extraction of tissues and phenotypes of tissue subpopulations using SIGN. (**A**) healthy tissue samples in GTEx, (**B**) tumor tissue samples in TCGA, (**C**) PDX tumor models in PDX encyclopedia, and (**D**) HSC and progenitors in hematopoietic hierarchy [32].

Moreover, we implemented SIGN on The Cancer Genome Atlas (TCGA) dataset of patient tumor samples [33] to identify the tumor type of each sample in the TCGA (Fig. 5B). Using leave-one-out cross validation, we could identify the tumor types, for the 24 populations with more than 70 samples (Supplementary Table 3), with 99.4% balanced accuracy. We explored the misidentifications in the tumor types with the lowest balanced accuracy such as Lung and Head and Neck cancer (Fig. 5B). We found out that some of the head and neck cancer (HNSC) patients are misidentified as lung cancer and some of the lung squamous cell carcinoma patients, in lung cancer cohort, are misidentified as HNSC which due to their similarity based on same cell type of origin, squamous cells [34].

Sampling tumors and growing them in animal models forces cells to evolve and adapt themselves to the microenvironment in animal models. Hence, we investigated if cells hold their tissue specificity at the transcriptomic level in patient-derived xenograft (PDX) models. In this regard, we implemented SIGN on RNA-Seq data of PDX models of 5 tumor types available in the recent Novartis PDX encyclopedia [35], which include non-small cell lung cancer (NSCLC), breast cancer (BRCA), colorectal cancer (CRC), pancreatic adenocarcinoma (PDAC), and cutaneous melanoma (CM) (Fig. 5C). Sensitivity of the predictions ranges from 82% (for NSCLC) to 100% (for CM).

### Identification of phenotypes in homogeneous population

We observed that SIGN yielded lower performance in identifying healthy and malignant tissue types with lower number of samples in the reference populations. We investigated if intra-sample heterogeneity among each population is a confounding factor in performance of SIGN. We implemented SIGN on RNA-Seq data of hematopoietic stem cells (HSC) and progenitors in hematopoietic hierarchy [36,37] as homogeneous populations. In spite of low sample number, 10 for HSC and ≤5 for progenitors, SIGN could identify samples in earliest thymic progenitors (ETP), common myeloid progenitors (CMP), granulocyte-monocyte progenitors (GMP), and megakaryocytic-erythroid progenitors (MEP) perfectly in spite of their similarity [32] (Fig. 5D). The identification of phenotypes in the hierarchy was significant in total (MCC=0.83, p<0.001). We observed 1 misidentification in multilymphoid progenitors (MLP) as multipotent progenitors (MPP), 2 MPP as HSC, and 2 HSC as MPP (Fig. 5D) which are in agreement with branching in hematopoietic hierarchy and high similarity of MPP and HSC phenotypes [36].

We explored pathways which were dissimilar between HSC and MPP as well as MEP and MLP as two pairs of highly similar and highly dissimilar phenotypes in hematopoietic hierarchy, respectively. We could identify many immune related hallmarks representative of differences between MEP and MLP populations such as *IL6 JAK STAT3 SIGNALING* and *INFLAMMATORY RESPONSE*. *REACTIVE OXYGEN SPECIES PATHWAY* as an oxygen metabolism hallmark was among the dissimilar hallmarks as well (Supplementary Fig. 1). These results are in agreement with the differences between MEP and MLP as progenitors responsible for erythrocyte and B/T cell productions, respectively [36,38]. Moreover, we could identify *WNT BETA CATENIN SIGNALING*, *NOTCH SIGNALING*, *TGF BETA SIGNALING*, and *CHOLESTEROL HOMEOSTASIS* as dissimilar hallmarks between HSC and MPP which agree with previous literature in this regard [39–43]

### Prediction of drug perturbations

Treating cancer cells with drugs may not necessarily result in changes in cell viability as cells may be resistant to therapies. However, they can affect signaling and metabolic pathways by changing expression of genes as components of biological pathways. Hence, we investigated if SIGN can classify drugs, based on their mechanism of action, which cancer cells have been exposed to in connectivity mapping (CMap) database [17] using the differences between expression of genes before and after treatment. We implemented SIGN on chemical perturbation for 7 classes of drugs including HDAC inhibitors, proteosome inhibitors, microtubule inhibitors, antipsychoics, Na+/K+ATPase pump inhibitors, HSP90 inhibitors, and CDK inhibitors (Supplementary Table 4). We could classify drugs in these classes significantly (MCC=0.94; p<0.001) (Fig. 6). Therefore, changes in gene expression patterns of biological pathways could be representative of the drug which the cancer cells have been treated with.

**Figure 6.**
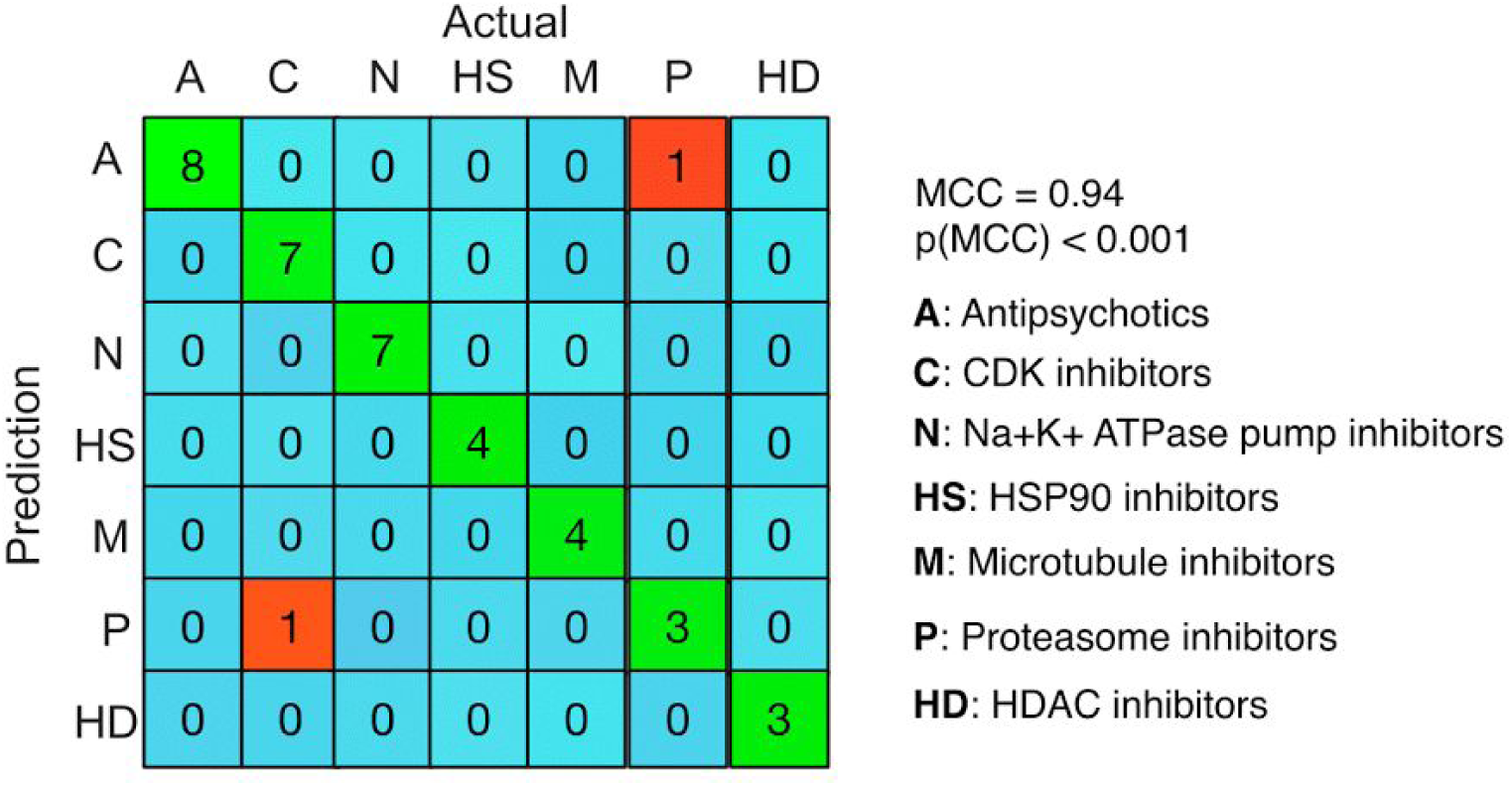
Confusion matrix of drug classification, based on their mechanism of action, by implementing SIGN on genetic perturbation data provided in connectivity mapping (CMap) [44] database.

## DISCUSSION

We introduced TSC, a new statistic capturing differences in expression patterns of biological pathways between biological samples from different conditions. We showed that this statistic overcomes the limitations of gene set and singular enrichment analysis methods [8] by considering variability of expression of genes within biological samples in a population. Moreover, it considers cooperative activity of genes within biological pathways. Hence, it also overcomes limitations of multivariate models which are not based on biological rationale behind choosing set of cooperative genes. We established performance of TSC by capturing oncogenic hallmarks for subtypes of breast cancer and showed stability of the results with respect to change in cohort of patients, imbalance in the compared populations (healthy versus tumor samples), as well as decreasing the number of tumor samples down to 10.

We used TSC to develop SIGN, a methodology for phenotype classification. This method uses distribution of TSC of biological pathways between a given sample and a set of references populations with known phenotypes to identify its phenotype same as the closest populations. In this approach, we do not limit the information space, such as expression of genes and their patterns in biological pathways, which makes the method less biased and more generalizable for different applications.

We illustrated the versatility and interpretability of SIGN as a new approach in identifying healthy and malignant biological samples. We stratified normal and tumor samples along with poor versus good prognosis patient samples in the largest gene expression dataset of breast tumors. Our results suggest that SIGN can be used for tumor diagnosis and prognosis. To investigate generalizability of SIGN, we investigated its performance in identifying the site of extraction of human tissue samples. More than 99% specificity in identifying site of extraction achieved for identification of 13 healthy and 24 tumorigenic site of extraction samples in GTEx and TCGA, respectively, showing specificity of gene expression pattern in each site in both healthy and malignant phenotypes. Although gene expression patterns will be changed upon engrafting tumors in patient derived xenografts, we showed that they still holds their specificity to each tumor type. Hence, SIGN can be used for identification of tumor type in patient-derived xenografts. Moreover, we clearly showed the high performance of SIGN in detecting samples using homogeneous populations with as low sample number as 3 by identifying phenotypes in hematopoietic hierarchy. In this regard, SIGN can be used to significantly identify homogeneous phenotypes or subpopulations within each tissue or tumor type.

Considering the dependency of states of cancer cells on the stress condition which the cells could have been exposed to, we showed that SIGN can be used to identify the type of stress based on perturbations in gene expression patterns. Using SIGN, we classified 7 classes of drugs based on their mechanism of action, along with the chemical perturbation data of the drugs provided in CMAP database. Hence, one can use SIGN to identify the classes of drugs which cancer cells (patient tumors) may have been exposed to.

Conclusively, SIGN is a generalizable classification method, considering its unbiased way of classifying biological samples, which can be applied to stratify patients based on their survival, classify biological samples relying on their site of extraction in vivo, and classifying types of drugs cancer cells have been exposed to. With the increasing amount of gene expression and transcriptomic data, we will investigate performance of SIGN in other applications such as identification of cancer stem versus non-stem cells, stratification of patients based on their survival across multiple cancer types, and identification of mechanism of action of new drugs using genetic perturbation data in CMAP database.

## ACKNOWLEDGEMENTS

The authors would like to thank Jacques Archambault for supporting Nehme El-Hachem financially. The figures of human organs in Fig. 2 have been extracted from at www.gettyimages.ca.

### AUTHOR CONTRIBUTIONS

S. A. M. T. defined TSC and developed SIGN. S. A. M. T. and V. S. K. M. performed the analysis and interpreted the results. S. A. M. T., V. S. K. M., and N.E.-H collected and curated public datasets for this study. S. A. M. T., V. S. K. M., and B.H-K conceived the design of the study. S. A. M. T., V. S. K. M., and B.H-K wrote the manuscript. B.H-K supervised the study.

### FUNDING

This study was conducted with the support of the Canadian Cancer Research Society and the Ontario Institute for Cancer Research through funding provided by the Government of Ontario. S. A. M. T was supported by Connaught International Scholarships for Doctoral Students. V. S. K. M. was supported by the Cancer REsearch Society. B.H.K was supported by the Gattuso-Slaight Personalized Cancer Medicine Fund at Princess Margaret Cancer Centre and the Canadian Institutes of Health Research.

### COMPETING FINANCIAL INTERESTS

The authors declare no competing financial interests.

